# FK506-binding protein 2 participates in proinsulin folding

**DOI:** 10.1101/2022.12.12.520056

**Authors:** Carolin Hoefner, Tenna Holgersen Bryde, Celina Pihl, Sylvia Naiga Tiedemann, Sophie Emilie Bresson, Hajira Ahmed Hotiana, Muhammad Saad Khilji, Theodore Dos Santos, Michele Puglia, Paola Pisano, Mariola Majewska, Julia Durzynska, Kristian Klindt, Justyna Klusek, Marcelo J. Perone, Robert Bucki, Per Mårten Hägglund, Pontus Gourdon, Kamil Gotfryd, Edyta Urbaniak, Malgorzata Borowiak, Michael Wierer, Patrick Edward MacDonald, Thomas Mandrup-Poulsen, Michal Tomasz Marzec

**Author notes:** Equal contribution (Carolin Hoefner, Tenna Holgersen Bryde, Celina Pihl). Corresponding author and person to whom reprint requests should be addressed:* Associate Professor Michal Tomasz Marzec M.D., PhD. Postal Address: Panum Institute 12.6.8, 3B Blegdamsvej, DK-2200 Copenhagen, Denmark.

## Abstract

Apart from chaperoning, disulphide bond formation and downstream processing, the molecular sequence of proinsulin folding is not completely understood. Proinsulin requires proline isomerization for correct folding. Since FK506-binding protein 2 (FKBP2) is an ER-resident proline isomerase, we hypothesized that FKBP2 contributes to proinsulin folding. We found that FKBP2 co-immunoprecipitated with proinsulin and its chaperone GRP94, and that inhibition of FKBP2 expression increased proinsulin turnover with reduced intracellular proinsulin and insulin levels. This phenotype was accompanied by an increased proinsulin secretion and the formation of proinsulin high molecular weight complexes, a sign of proinsulin misfolding. FKBP2 knockout in pancreatic β-cells increased apoptosis without detectable upregulation of ER-stress response genes. Interestingly, FKBP2 mRNA was overexpressed in β-cells from pancreatic islets of T2D patients. Based on molecular modelling and an in vitro enzymatic assay, we suggest that proline at position 28 of the proinsulin B chain (P28) is the substrate of FKBP2’s isomerization activity. We propose that this isomerization step catalyzed by FKBP2 is an essential sequence required for correct proinsulin folding.

## Introduction

Misfolding of the insulin prohormone, proinsulin (PI), is increasingly appreciated as an early pathogenic event in type 2 diabetes (T2D) when a mismatch is reached between PI biosynthetic demand, typically increased due to insulin resistance, and folding capacity [1, 2]. PI misfolding perturbs the insulin production machinery causing decompensation of insulin secretion relative to needs and is believed to eventually contribute to a reduction of functional β-cell mass due to ER stress-induced apoptosis [1–4]. PI folding involves multiple steps, including chaperoning and disulphide bridge formation [5, 6], all facilitating the locking of the linear amino acid chain into the 3D structure that after further processing to mature insulin lends the biological activity to the molecule.

The amino acid proline (Pro) adopts either a *cis* or *trans* isomeric conformation as it is incorporated into the polypeptide chains. The relatively low energy difference between *cis* and *trans* isomers of any amino acid (X) to Pro peptide bonds typically allows the presence of 3-10 % *cis* X-Pro isomers in proteins, depending on the upstream amino acid. Nonetheless, usually only *trans* conformation is favorable for protein folding [7, 8].

The transition of a proline residue from *cis* to *trans* conformation, or vice versa, is called *cis-trans* isomerization. This process occurs within the nascent amino acid chain and is a rate-limiting early step in protein folding that can trap a proline residue in an isomeric state, preventing proper folding [9, 10]. Human PI contains three prolines, of which only the proline at position 28 (P28) in the B chain is evolutionarily conserved (see below). The only publicly available PI crystal structure of folded PI contains an amino acid swop at the P28 position (P28K29 to K28P29) and molecule P29 is in *trans* conformation (Protein Data Bank structure [11]). During protein synthesis, the majority of P28 residues are expected to be *trans*-proline isomers and to fold readily. The remaining P28 residues will be *cis*-proline isomers and will require *cis-to-trans* isomerization to allow subsequent folding.

Proline isomerization is catalyzed by peptidyl prolyl *cis-trans* isomerases [12–14]. Interestingly, broad pharmacological inhibition of these isomerases with e.g. the potent immunosuppressant FK506 deteriorates insulin mRNA levels, glucose-stimulated insulin secretion, β-cell survival and leads to diabetes development in immunosuppressed patients posttransplantation [15–19]. Recently a P28 to leucine mutation has been reported in a case of maturity-onset diabetes of the young (MODY) presenting defects in PI folding, trafficking, and dominant-negative behavior in vitro, emphasizing the clinical importance of *cis-trans* isomerization in insulin biosynthesis [20].

FK506 binding protein 2 (FKBP2) is one of seven proline isomerases expressed in the ER, where it partakes in protein folding and is induced in response to the buildup of unfolded proteins during ER stress [21, 22]. Despite this, the substrates of FKBP2 are yet to be determined. Taken together, we hypothesized that P28 isomerization is an early event during PI folding executed by FKBP2.

Here we show that FKBP2 is a novel PI binding partner in model β-cells and that P28 is a target for FKBP2-dependent isomerization. We demonstrate that FKBP2 binds to the recently described PI chaperone Glucose Regulated Protein 94 (GRP94) which is essential for PI handling [5]. Additionally, we observed that FKBP2 knock out (KO) in insulin-producing cells leads to loss of intracellular PI via increased misfolding, loss of PI solubility and increased secretion of immature PI. Furthermore, while ER stress markers remain unchanged, FKBP2 KO results in increased cellular apoptosis. Finally, we show that FKBP2 mRNA is overexpressed in β-cells in human islets from organ donors with T2D patients, likely as a compensatory response.

## Research Design

### Cell culture

Rat insulinoma INS-1E cells (wild-type, FKBP2 control and FKBP2 KO) were cultured in RPMI-1640 complete medium (for details, see supplementary procedures) at 37°C with 5 % CO_2_.

### Generation of FKBP2 CRISPR/Cas-9 mediated knockout INS-1E cell lines

FKBP2 knockout INS-1E cells were generated with ready-to-use Lentiviral plasmids encoding a guide-RNA (gRNA) sequence targeting rat *fkbp2* exon 2 (supplementary procedures).

### FKBP2 expression in pancreatic endocrine cells

The raw sequencing reads on human non-diabetic islets (deposited at EMBL-EBI under accession number E-MTAB-5061) and processed gene expression matrices along with the cell type inferred, were reanalyzed searching for FKBP2 mRNA presence in human adult islets, including β-cells. Human pancreas sections were obtained from Collegium Medicum in Bydgoszcz or Nicolaus Copernicus University in Torun (Poland) under protocols approved by the Bioethical Commission (KB381/2020). Human pancreata were dissected from 3 non-diabetic donors: 45-year-old male (BMI 22.3, A1c 4.9 %), 59-year-old female (BMI 35.2, A1c 6.0 %), and 62-year-old female (BMI 29.6, A1c 5.2%). Paraffin section (four μm) sections using standard protocol were prepared. Three to five sections per donor were stained overnight with primary antibodies (Suppl. Table 2) followed by secondary antibodies conjugated with AlexaFluor488, TRITC or AlexaFluor647 (Jackson Immunoresearch, USA) prior to imaging. Adjacent tissue sections were stained with hematoxylin-eosin and evaluated by pathologist for the absence of abnormal cellular phenotypes (supplementary procedures).

### Immunoprecipitations and mass spectrometry analysis

INS-1E cells were transfected at 60 % confluence using Lipofectamine™ 3000 (ThermoFisher Scientific, Denmark) with plasmids coding for GFP-tagged GRP94, PI, ER-localized GFP [23, 24] or myc-tagged FKBP2, according to the manufacturer’s protocol. Proteins were immunoprecipitated (IP) using the magnetic GFP- or myc-trap beads (Chromotek, Germany). IP samples were reduced and alkylated, digested with trypsin/LysC, and the resulting peptides were analyzed on a Bruker Impact II ESI-QTOF (Bruker Daltonics, USA) mass spectrometer (supplementary procedures).

### Size Exclusion Chromatography

Whole-cell lysates were fractionated by size exclusion chromatography (SEC) using a SuperdexTM 75 10/300 GL column (SigmaAldrich, Denmark) in Tris buffer (50 mM Tris pH 8.0, 150 mM NaCl, 5 mM KCl). Fractions of interest were analyzed via SDS-PAGE and immunoblotting (supplementary procedures).

### Immunoblotting

Proteins were detected in cellular lysates, soluble and non-soluble (pellets) fractions and cell culture supernatants using immunoblotting with specific antibodies (Suppl. Table 3) under non-reducing and reducing conditions (supplementary procedures).

### Single-cell RNA sequencing

Published datasets of single-cell RNA sequencing (scRNAseq) from dual patch-clamp sequencing studies were used to evaluate FKBP2 expression [25, 26]. Cell types were assigned by Uniform Manifold Approximation and Projection with Leiden Clustering, and the expression of insulin, glucagon, pancreatic polypeptide, or somatostatin was analyzed.

### *In silico* modelling of protein interactions and protein sequences analysis

Using the ZDOCK 3.0.2 modelling software [27], crystal structures from the protein data bank [28] of FKBP2 (PDB ID: 2PBC [29]) and PI (PDB ID: 2KQP [11]) were used to generate prediction models of their interaction. Using the Python script ‘InterfaceResidues’ [30], residues predicted to interact were identified in PyMOL (Schrödinger, USA), using a 2.5 Å vicinity as a cut-off. FKBP2 evolutionary conserved residues were identified based on literature [31] and alignment of protein sequences among FKBP family members. FKBP2 promotor analysis was done within 500 bp upstream of the FKBP2 startsite using TRANSFAC 2.0. Insulin 1/2 and IGF 1/2 amino acid sequences were retrieved from UniProtKB v. 2021_04 and aligned with BlastP.

### FKBP2 protein activity assay

n-Nitroanilide modified peptide (GERGFFYTPF-F-pNA, GenScript, USA) was mixed with assay buffer (DPBS pH 7.4), recombinant FKBP2 (1 μg, Abcam, Great Britain) or assay buffer alone and/or chymotrypsin (20 mM, Roche, Switzerland) to initiate the reaction. The increase in absorbance (405 nm) was read in 1 minute intervals for 12 hours at 25°C. The signal from the time point with the highest absorbance in the reaction that contained FKBP2, n-Nitroanilide modified peptide and chymotrypsin was used to compare experimental conditions and generate the graph (supplementary procedures).

### qRT-PCR

The relative mRNA level of ER stress markers and insulin genes was determined by quantitative RT-PCR using specific primers (Suppl. Table 4; supplementary procedures).

### Apoptosis Assay

Apoptotic cell death was determined by the detection of DNA–histone complexes present in the cytoplasmic fraction of cells using the Cell Death Detection ELISAPLUS kit (Roche, Switzerland) according to the manufacturer’s protocol (supplementary procedures).

### Glucose-stimulated insulin-secretion (GSIS)

INS-1E cell lines (control, FKBP2 KO) were examined for PI and insulin secretion in response to 2 and 20 mM glucose according to standard protocols (supplementary procedures).

### Statistics

Differences between the two groups were assessed by two-tailed Student’s t-test, or by one-way ANOVA of treatments versus control using the GraphPad Prism 9 (La Jolla, USA). Data are presented as means± SD. P-values ≤ 0.05 were considered significant.

## RESULTS

### FKBP2 directly interacts with proinsulin and FKBP2 mRNA is overexpressed in islets from type 2 diabetes patients

To identify new PI interacting partners with the potential to impact PI folding we exogenously expressed GFP-tagged PI and GRP94 (a novel PI chaperone [5]) in INS-1E cells. The GFP-tag enabled us to immunoprecipitate substantial amounts of target proteins for the detection and identification of co-immunoprecipitated partners during subsequent mass spectrometry analysis (Fig. 1A). Among proteins localizing to the ER lumen, 11 were found to bind both PI and GRP94 (Fig. 1A), including FKBP2 (supp. Fig. 1). Western blotting (WB) analysis confirmed the co-precipitation of Myc-tagged FKBP2 to PI and GRP94 (Fig. 1A).

**Fig. 1.**
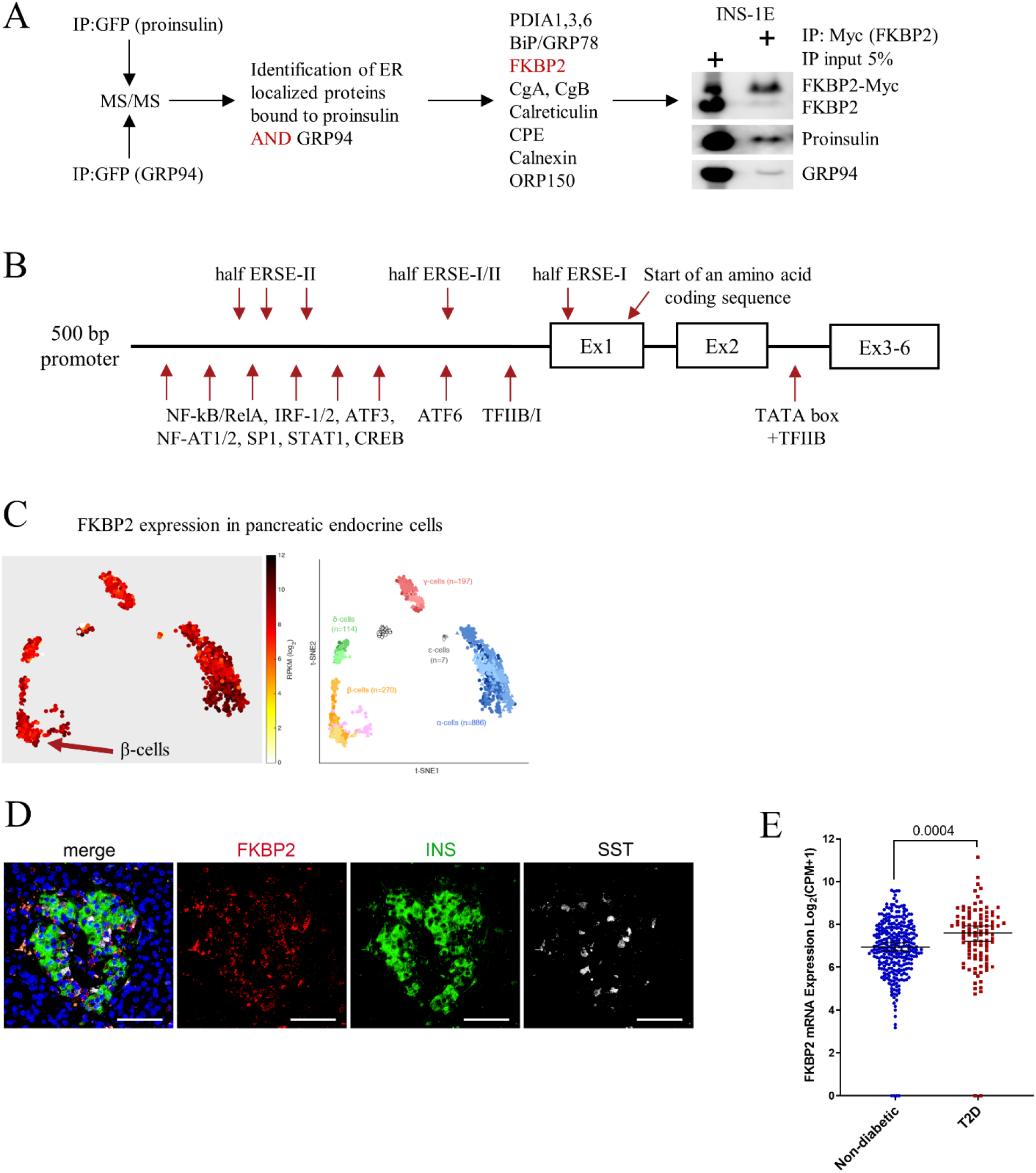
FKBP2 isomerase interacts with proinsulin and GRP94 and is expressed in β-cells. **A.** INS1-E cells were transiently transfected to express GFP-tagged human proinsulin and GRP94, both of which were immunoprecipitated and analysed by mass spectrometry for binding partners present in the endoplasmic reticulum. FKBP2 interaction with proinsulin and GRP94 was confirmed by SDS-PAGE/Western blot analysis of the immunoprecipitates of myc-tagged FKBP2 expressed in INS1-E cells. PDIA 1, 3, 6: Protein disulfide-isomerase 1, 3, 6; BiP/GRP78: Binding immunoglobulin protein/ Glucose Regulated Protein 78; CgA, CgB: Chromogranin A and B; CPE: Carboxypeptidase E; ORP150: Oxygen-regulated protein 150. **B.** Graphic representation of FKBP2 promoter, with putative transcription factors binding sites and regulatory elements identified using PROMO from ALGGEN. Analysed DNA sequence was retrieved from HGNC:3718. ERSE: ER stress response element; TFIIB/I: Transcription factor IIB/I; ATF6 and 3: Activating transcription factor 6 and 3; NF-kB/RelA Nuclear factor-kB/REL-associated protein; IRF-1/2: Interferon regulatory factor 1/2; NF-AT1/2: nuclear factor of activated T cells; SP1: Specificity Protein 1; STAT1: Signal transducer and activator of transcription 1; CREB: cAMP Response Element-Binding Protein; Ex: exon. **C.** FKBP2 single-cell gene expression. t-SNE representations coloured according to FKBP2 expression levels, n=1554 from Segerstolpe et al, Cell Metabolism 2016. **D.** Representative fluorescence microscopy images of the human adult pancreas stained with antibodies against FKBP2 (red), INS (insulin, green) labeling β-cells and SOMATOSTATIN (SST) (white) labeling δ-cells. Scale bars = 100 μm. DAPI (blue) labels the nuclei. **E.** *FKBP2* mRNA expression analyzed by single cells RNA sequencing of β-cells derived from healthy and T2D individuals. Lines represent median with 95% CI error bars. Statistical analysis was performed using unpaired two-tailed parametric t-tests. FKBP2 transcript expression is shown as log2 (counts per million + 1).

The murine FKBP2 promoter contains regulatory elements characteristic for genes induced during the unfolded protein response (UPR) [32], and analysis of the 5’ flanking region of human *FKBP2* identified similar regulatory elements e.g. SP1, ATF3/6 and 3 incomplete ER stress response element sites (ERSE, Fig. 1B), suggesting a role for FKBP2 in maintaining ER homeostasis.

Single-cell *FKBP2* gene expression analysis revealed *FKBP2* mRNA in all pancreatic islet cell types (Fig. 1C, data reanalysis from [33]), and FKBP2 protein expression was confirmed in human β-cells (Fig. 1D). As the increased demand for insulin in T2D may require compensatory up-regulation of chaperones involved in PI folding, we expanded the analysis to T2D cases. The results showed a significant up-regulation of *FKBP2* in β-cells of T2D donors (Fig. 1E, [25, 26]), however, these data have not been independently validated (e.g. by qPCR).

### Proinsulin P28 is a potential target of FKBP2-dependent isomerization

Through ZDOCK modelling software and crystal structures of FKBP2 (PDB ID: 2PBC) and PI (PDB ID: 2KQP), their interaction was modelled, demonstrating that PI fits into the FKBP2 substrate binding pocket (Fig. 2A-B). Of note, FKBP2 exists mainly as a monomer in the β-cell, as evidenced by size exclusion chromatography (SEC) analysis (Supp. Fig. 2). The majority of PI residues interacting with FKBP2 are located in the PI B-chain (Fig. 2C) and include P28. On FKBP2’s side of the interaction, almost all evolutionary conserved residues presumed to facilitate FKBP2’s isomerase activity interact with PI (Fig. 2D). The PI 2KQP structure (Fig. 2B, C) represents PI with an introduced mutation swopping the P28-K29 positions. Despite this, the *in silico* analysis identified P28 (P29 in the crystal structure) as the only plausible site of FKBP2-dependent proline isomerization. Furthermore, the P28 residue is fully evolutionary conserved within PI and the Insulinlike growth factor family of hormones (Fig. 2E-F), underscoring its structural and/or functional significance.

**Fig. 2.**
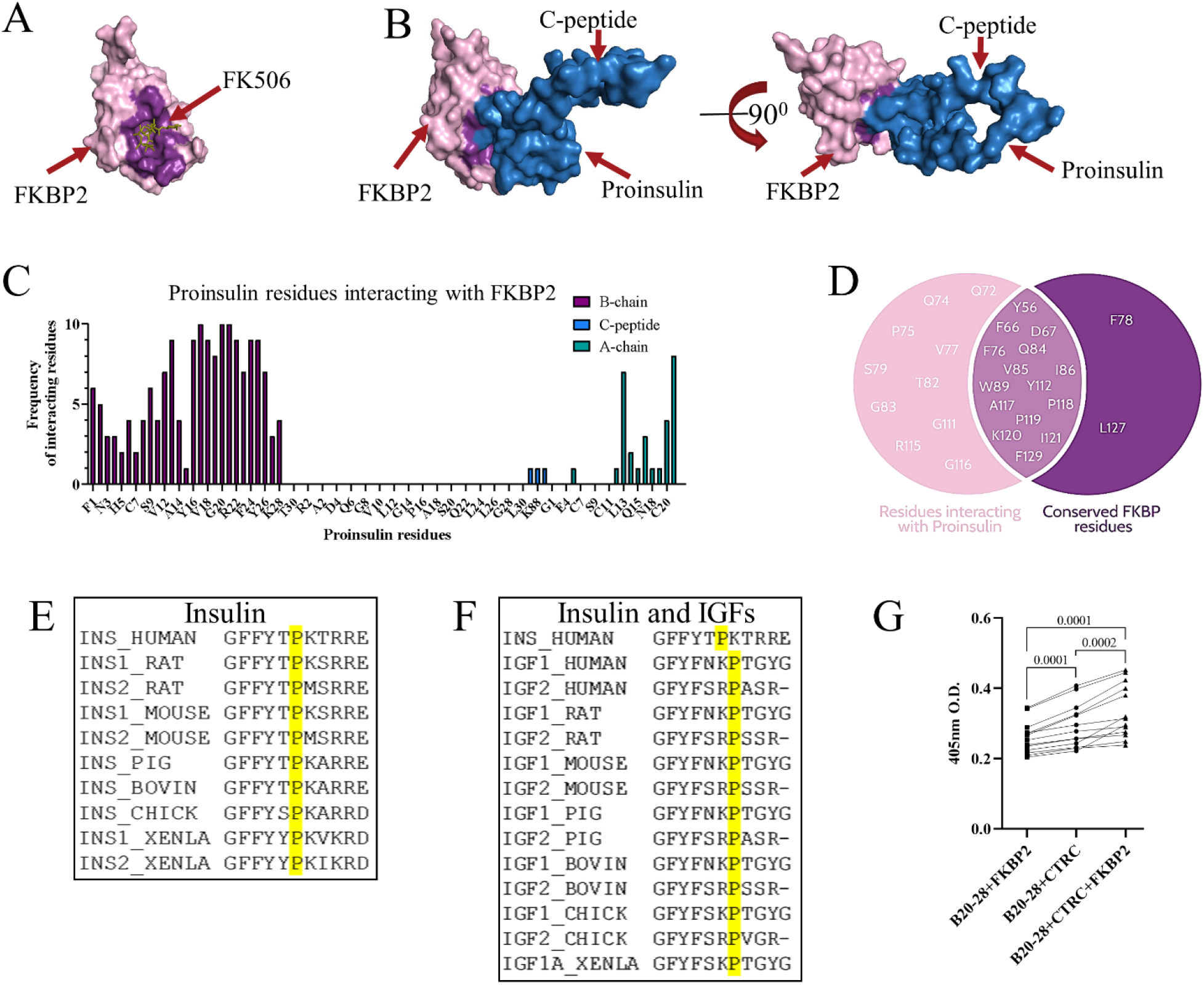
In silico modelling of proinsulin interaction with FKBP2, identification of proinsulin isomers and demonstration of FKBP2 as a proinsulin isomerase. FKBP2 binding to its inhibitor FK506 (**A**, crystal structure PDB: 4NNR) and proinsulin (**B**, FKBP2 crystal structure PDB: 2PBC, proinsulin crystal structure PDB: 2KQP, blue) in the representative *in silico* model. In both structures, the residues involved in the interaction are highlighted in purple. The models were generated on the ZDOCK server. **C.** The panel shows the frequency of proinsulin residues interacting with FKBP2 in the top 10 generated interaction models with FKBP2 and proinsulin (2PBC and 2KQP crystal structures respectively). **D.** Overlap between proinsulin interacting residues and conserved residues within FKBP2. The data displayed are the combined data from 100 FKBP2-proinsulin interaction models. The conserved residues are highlighted in dark purple. **E-F.** Alignment of human proinsulin sequence (UniProt P01308), between positions 23 of B chain and position 1 of C peptide with proinsulins (**E**) and IGF-1/2 (**F**) of Homo sapiens (HUMAN IGF1 P05019, IGF2 P01344), Rattus norvegicus (RAT INS1 P01322, INS2 P01323, IGF1 P08025, IGF2 P01346), Mus musculus (MOUSE INS1 P01325, INS2 P01326, IGF1 P05017, IGF2 P09535), Sus scrofa (PIG INS P01315, IGF1 P16545, IGF2 P23695), Bos Taurus (BOVIN INS P01317, IGF1 P07455, IGF2 P07456), Gallus gallus (CHICK INS P67970, IGF1 P18254, IGF2 P33717) and Xenopus laevis (XENLA INS1 P12706, INS2 P12707, IGF1A P16501). Sequences were retrieved from UniProt v. 2021_04. **G.** Peptidyl-prolyl cis-trans isomerisation assay of peptide containing Pro28 (GERGFFYTP-F-pNA, B20-28) incubated with or without FKBP2. The substrate peptide was dissolved in LiCl/TFE buffer, and to initiate the reaction, chymotrypsin (CTRC) was added to digest and release the fluorophore (pNA) from the peptide in trans conformation. The absorbance was acquired at 405 nm. The signal from the time point with the highest absorbance in the reaction that contained FKBP2, n-Nitroanilide modified peptide and chymotrypsin was used to generate the graph. Data were analyzed by paired t-test between individual tested conditions.

Furthermore, we set up a proline *cis-trans* isomerization assay, where a peptide containing PI B chain residues 20-28 was modified to have the last residue, P28, followed by phenylalanine (F) linked to the fluorophore paranitroanilin (pNA). The peptide bond between P28-F-pNA can be proteolyzed by chymotrypsin only when proline is in *trans* conformation [34]. The peptide was synthesized using *cis* and *trans* prolines and thus should contain a mixture of isomers. The highest absorbance, representing free fluorophore pNA was observed when FKBP2 was added to the reaction (Fig. 2G), indicating that FKBP2 increased the presence of *trans* prolines in the reaction mixture.

### Diminished intracellular proinsulin and insulin contents after FKBP2 KO

We performed CRISPR/Cas9 induced KO to evaluate the effects of FKBP2 deficiency in INS-1E cells on PI and intracellular insulin levels (Supp. Fig. 3). FKBP2 KO cells cultured for 2 h in 20 or 2 mM glucose-containing media showed a significant reduction in intracellular PI contents (40 % compared to control) as evidenced by WB analysis (Fig. 3A). Similarly, insulin levels were reduced (40 %) at 20 mM glucose conditions. We observed no significant change in insulin levels at 2 mM conditions (Fig. 3A). Reconstitution of FKBP2 expression led to an increase in intracellular PI levels, but had no effect on insulin contents at 20 mM glucose (Fig. 3B).

**Fig. 3.**
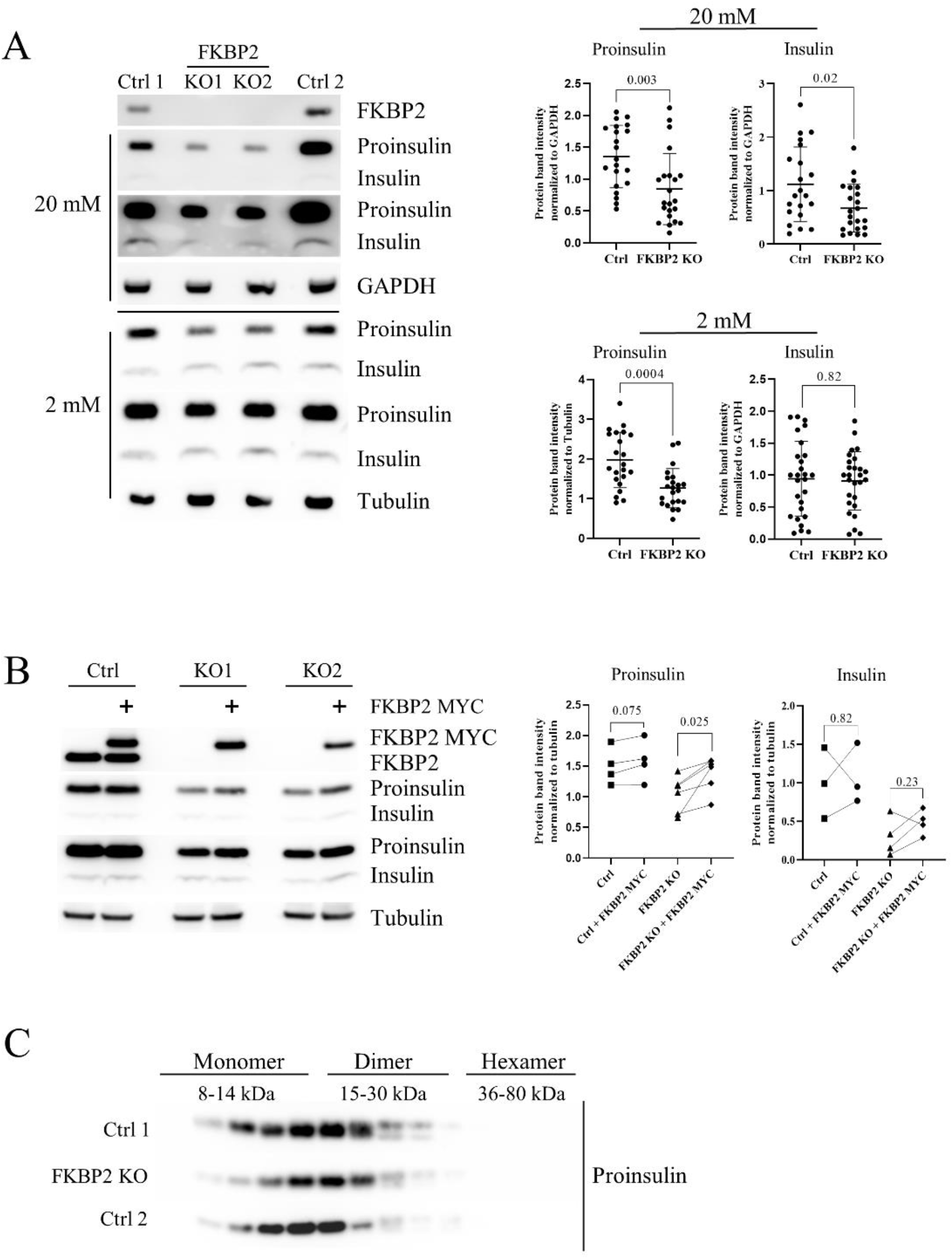
Diminished intracellular proinsulin and insulin contents after FKBP2 KO. **A.** SDS-PAGE and Western blot analysis of proinsulin and insulin expression levels in FKBP2 KO and FKBP2 WT cells (failed FKBP2 KO and INS1-E) after 2 h in 20 or 2 mM glucose containing media. FKBP2 KO was achieved via CRISPR/Cas9 guide FKBP2 directed-RNA targeting (clonal cell lines shown 3-6 months after viral transduction; representative blots of n>10 on the left and band quantification on the right). Proinsulin and insulin were visualized with ant-insulin antibody from Cell Signaling. **B.** SDS-PAGE and Western blot analysis of proinsulin and insulin expression levels upon exogenous expression of FKBP2 in FKBP2 KO clones and INS1-E (Ctrl) cells. Cells were transfected with plasmids coding for myc-tagged FKBP2, cultured for 48h and lysed 2 h after introduction of fresh culture media supplemented with 20 mM glucose. Representative blot of n=4 presented on the left and band quantification on the right. **C.** SDS-PAGE and Western blot analysis of proinsulin from FKBP2 KO, FKBP2 WT (Ctrl 1) and INS1-E (Ctrl 2) cells after size exclusion chromatography (SEC) separation at the pH 7,4 condition of cell lysis and SEC separation. SEC experiments n=3. Data for A-B represent means±SD analyzed by non-paired (A) or paired (B) t-test of treatments versus control.

Mutations resulting in the swop of PI residues P28 and K29 create conformational alterations that inhibit or weaken the formation of insulin dimers, allowing for more rapid absorption of fast-acting pharmaceutical insulins e.g. LisPro [35, 36]. Moreover, the amino acid sequence of pro-IGF-1, which does not naturally dimerize (but undergoes oligomerization [37]), resembles that of non-dimerizing insulins (Fig. 2F). Considering this along with FKBP2-PI modelling indicating P28 as a site of isomerization, we performed SEC separation of intracellular PI based on molecular mass. We detected no difference in the presence of PI monomers and dimers in FKBP2 KO cells when compared to the controls (Fig. 3C), indicating that potential FKBP2-dependent P28 isomerization does not influence PI dimer formation.

### FKBP2 KO does not lead to ER stress but induces β-cell apoptosis

We then examined the activation of the UPR pathways in the FKBP2 KO cells. We investigated mRNA expression levels of *BiP, PERK, CHOP, ATF6, IRE1* and unspliced and spliced levels of *XBP-1* and observed no statistically significant upregulation in FKBP2 KO cells when compared to controls. Eighteen h of treatment with 0.7 μM thapsigargin served as a positive control and induced a robust ER stress response (Fig. 4A). ER stress has been reported to induce *INS-1/2* mRNA degradation by activating the endonuclease IRE-1 [38–40], yet the results indicated that Ins-1/2 mRNA levels were not significantly reduced in FKBP2 KO cells (Fig. 4A), supporting our conclusion that this ER stress activated pathway is not triggered. We reasoned that FKBP2 KO and the loss of intracellular PI might induce cell apoptosis by means independent of ER stress (discussed below). Using a DNA–histone complex detection assay, we observed a significant increase in apoptosis in FKBP2 KO cells compared to controls, which was further upregulated upon thapsigargin treatment (Fig. 4B).

**Fig. 4.**
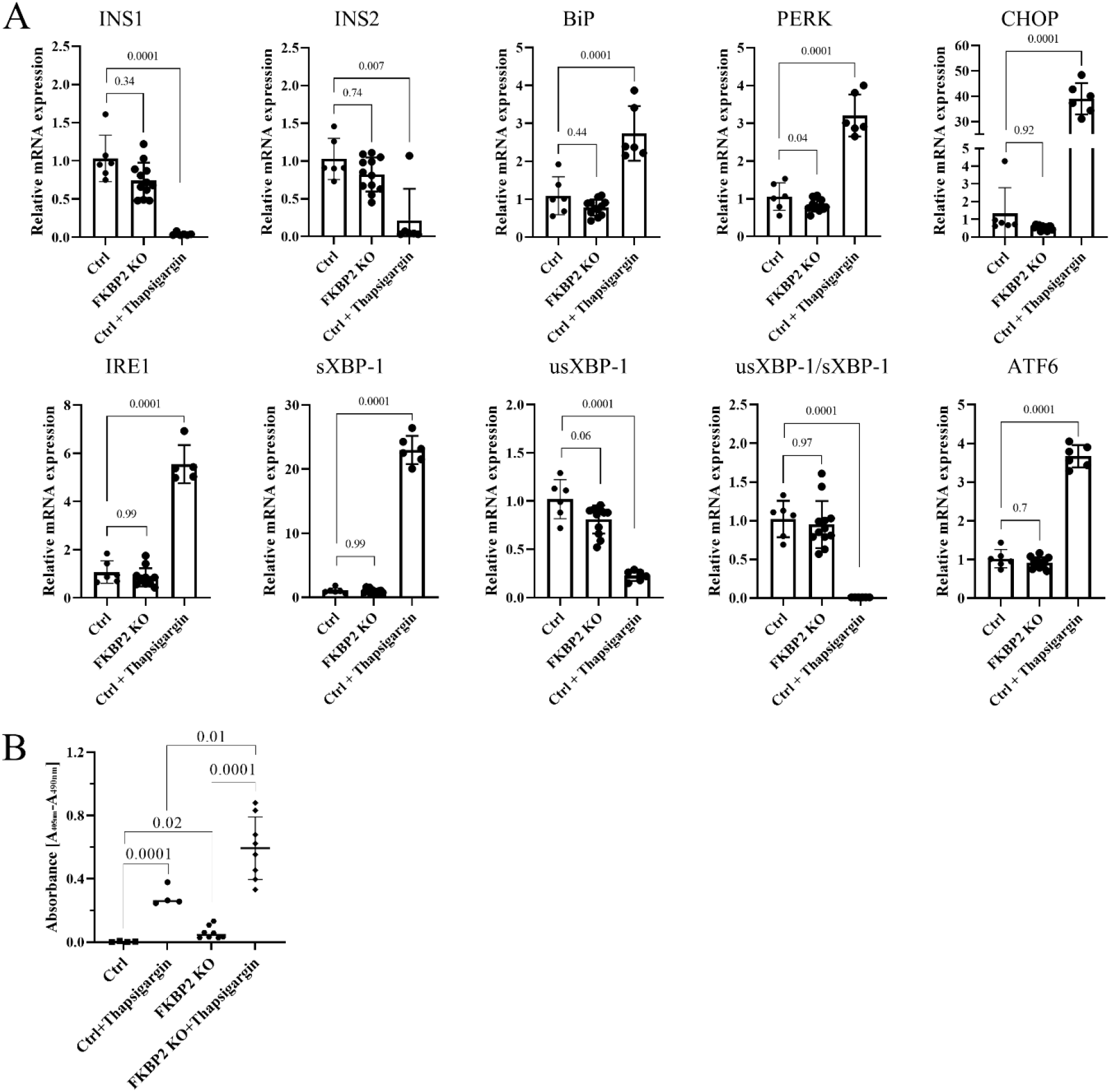
FKBP2 knockout does not induce endoplasmic reticulum stress but sensitizes cells to apoptosis. **A.** mRNA levels of genes in the endoplasmic reticulum stress pathways were analyzed by quantitative reverse transcription-PCR (qRT-PCR) in Ctrl (FKBP2 WT cells) and FKBP2 KO cells. Data represent the mean±SD analyzed by ordinary one-way ANOVA of treatments versus control, n>6. **B.** Apoptosis levels, representing internucleosomal degradation of genomic DNA, were analyzed in FKBP2 KO and Ctrl cells, n>4. Where indicated, cells in A and B were exposed for 18 h to 0,7 μM of thapsigargin.

### Proinsulin turnover is increased in FKBP2 KO cells accompanied by increased proinsulin secretion

As *INS-1/2* mRNA levels were comparable between control and FKBP2 KO cells (Fig. 4A), but steady-state PI levels were lower in FKBP2 KO cells, we reasoned that PI degradation at the ER stage or in the post-ER compartments during FKBP2 deficiency results in increased intracellular PI protein turnover. Consistently, following inhibition of protein synthesis with 100 μM cycloheximide (CHX), cellular PI levels were reduced by 50 % after 2 hours in FKBP2 KO lysates, whereas only 20 % reduction was observed after 4 hours in control cell lysates (Fig. 5A-B), demonstrating shorter PI protein half-life in FKBP2 KO cells. Brefeldin A (BFA) treatment has been shown to efficiently block ER to Golgi protein translocation and thereby inhibit PI secretion [6]. To determine if the shorter half-life was caused by an increased translocation and secretion, we pre-treated cells with 200 nM of BFA inhibiting ER-Golgi anterograde transport before cycloheximide exposure. BFA did not change PI half-life significantly between Ctrl and Ctrl+BFA or FKBP2 KO and FKBP2 KO+BFA. However, when those changes were analysed between pooled Ctrl (CHX and CHX+BFA) and FKBP2 KO (CHX and CHX+BFA) conditionss, the average change of PI content over the time course of the experiment was reduced in the Ctrl group and increased in the FKBP2 KO and the pattern was statistically significant (Fig. 5B bottom). Taken together, this result could indicate that our observations are the result of a combined cytotoxic effect of BFA (lowering PI content) and primary inhibition of PI secretion (increase in PI content), and suggest that PI secretion plays a role in shortening PI half-life in the FKBP2 KO phenotype.

**Fig. 5.**
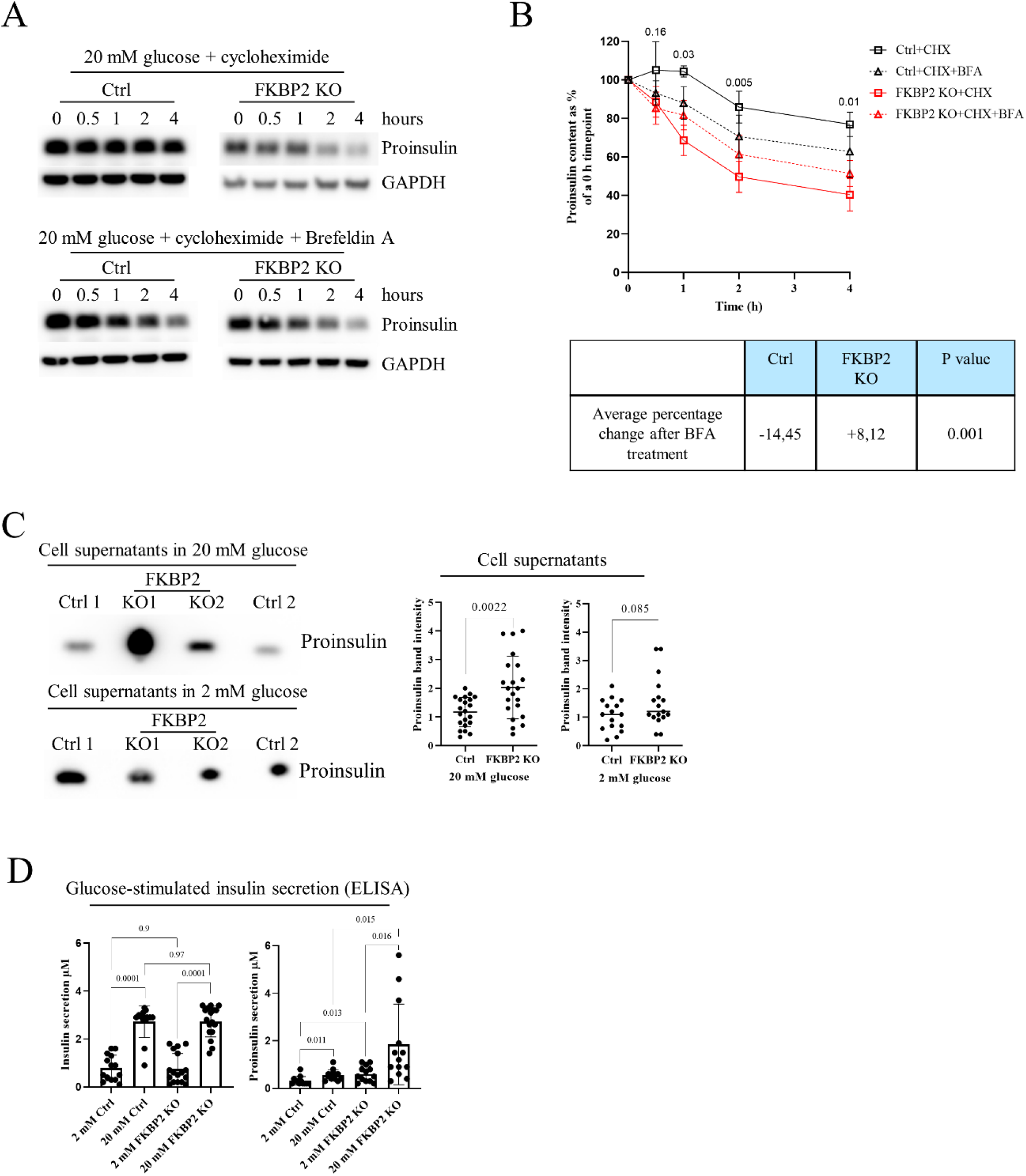
FKBP2 KO shortens proinsulin intracellular half-life and increases it secretion. **A-B.** INS-1E control (Ctrl) and FKBP2 KO cells were cultured for 3 h in 2 mM glucose-containing media, and subsequently 100 μM of the protein synthesis inhibitor cycloheximide (CHX, upper panels) and 200 nM of the inhibitor of exocytosis Brefeldin A (BFA, bottom panels) was added to the culture media from the start of the experiment. Cells were lysed at indicated time points, analyzed via reducing SDS-PAGE and proinsulin and insulin visualized through Western blotting (WB) with anti-pro-insulin antibody, n=5. B. WB data quantification. Top: graph p values were calculated by unpaired t-test of Ctrl+CHX vs FKBP2 KO+CHX for each time point. Bottom: Table with averages of changes of Ctrl and FKBP2 KO cells over the course of experiment in proinsulin content (band intensity) in response to BFA treatment. **C.** Accumulated secretion of proinsulin over the period of 6 h in 2 or 20 mM glucose by failed FKBP2 KO (ctrl 1 and 2) and FKBP2 KO cells, analyzed by SDS-PAGE and WB (n=10). 500 μL of cell supernatants (adjusted to cell number) was concentrated using 10 kDa MWCO filters to remove salts and reduce volume to 15 μL. Quantification of proinsulin bands was carried out with ImageJ (A and B) and normalized to GADPH (A) bands. Data are presented as means±SD analyzed by unpaired t-test of treatments versus control. **D.** The same cell types were tested for their ability to secrete insulin in response to the given glucose concentrations in KRBH buffer for a period of 30 min. Supernatants were analyzed by ELISAs specifically detecting mature insulin (left graph) or proinsulin (right graph) only. The bars represent the means ± SD.

To further examine this, we analysed PI secretion during a 4 h 20 mM glucose-stimulation and found that FKBP2 KO cells secrete significantly more PI than control cells, with this difference being less pronounced at 2 mM glucose stimulation, showing only a statistical trend (Fig. 5C), further suggesting that PI secretion is at least partially responsible for the observed reduction in intracellular PI levels.

Finally, we expected that FKBP2 KO-related changes in PI folding would lead to a lower secretion of insulin during glucose-stimulated insulin secretion (GSIS). However, that was not the case as no observable difference in mature insulin secretion in response to glucose stimulation was reported comparing control and FKBP2 KO cells (Fig. 5D). Interestingly, the same experimental supernatants showed increased PI secretion at 2 and 20 mM glucose conditions (Fig. 5D), aligning with the results from accumulated PI secretion (Fig. 5C).

### FKBP2 KO induces proinsulin misfolding and decreases its solubility

As FKBP2 KO induces intracellular PI loss, we wanted to identify the stage of PI synthesis involving FKBP2. So far, disulphide bond formation is the only discrete step in PI folding described [41]. To evaluate if FKBP2 preferentially binds to unfolded, reduced PI or oxidized, folded PI, we pretreated INS-1E cell lysates for 10 min with 100 mM of reducing agent 2-mercaptoethanol (2-ME) prior to FKBP2 immunoprecipitation. We observed a substantial increase in FKBP2 binding to reduced PI i.e. before disulfide bond bridges are formed (Fig. 6A), suggesting that PI-FKBP2 interaction occurs at an early PI folding stage. Furthermore, we evaluated the presence of PI high molecular weight complexes (PI multimers), as they represent faulty disulphide bond formation and are indicative of PI misfolding ([42] and Suppl. Fig. 4). When analyzed by non-reducing SDS-PAGE, control and FKBP2 KO cells showed similar levels of PI multimers at 2 and 20 mM glucose concentrations (Fig. 6 B). However, the proportion of monomers/multimers in FKBP2 KO was significantly decreased, due to lower levels of PI monomers detected after treatment with at 20 mM glucose.

**Fig. 6.**
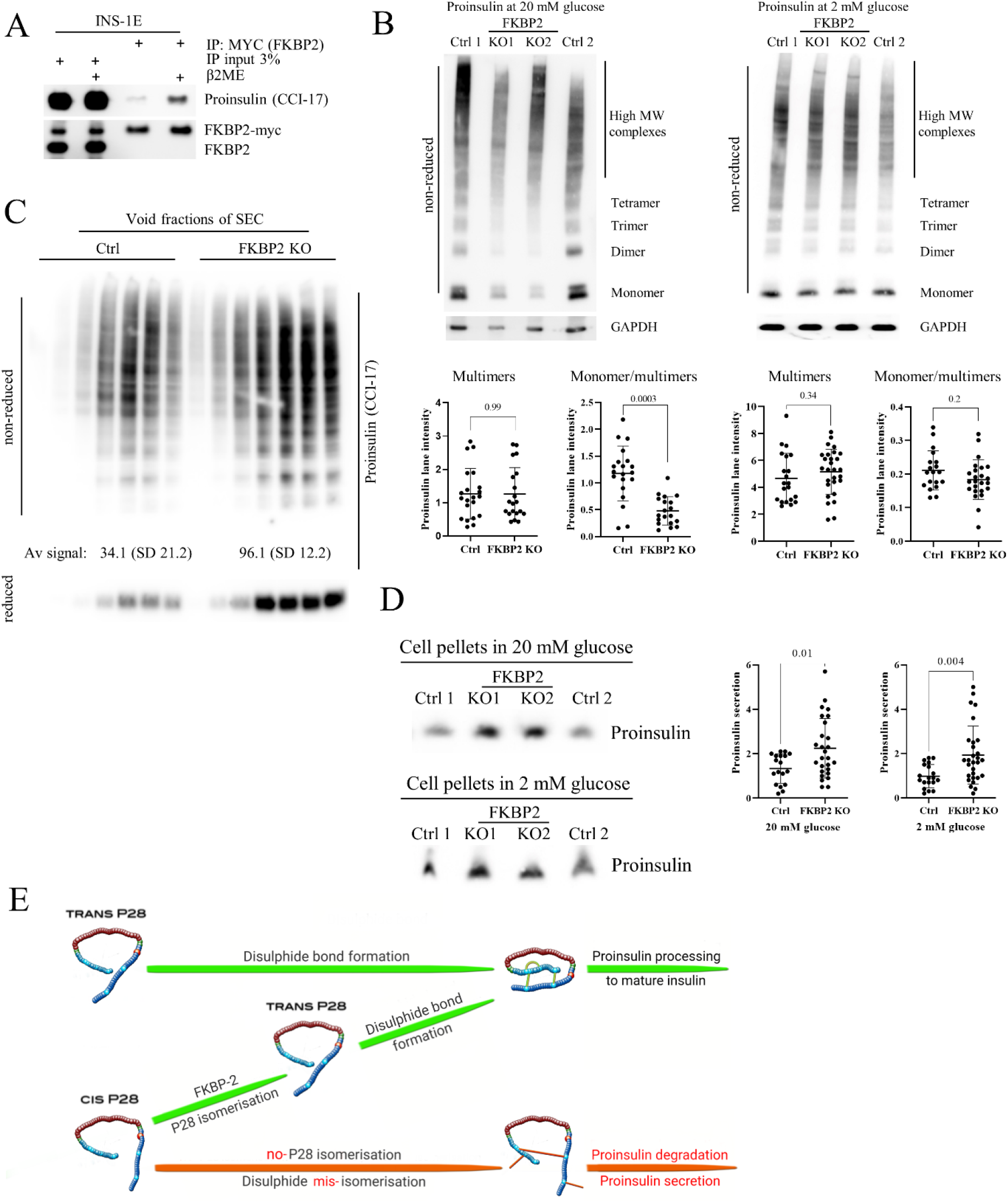
FKBP2 KO increases intracellular levels of high molecular weight proinsulin complexes and non-soluble proinsulin fraction. **A.** INS-1E cells were transfected with myc-tagged FKBP2, followed 48 h later with immunoprecipitation (via myc tag) and SDS-PAGE/Western blotting to detect proinsulin. Where indicated, cell lysates were pretreated prior to immunoprecipitation for 10 minutes with 100 mM of reducing agent β2 mercaptoethanol (β2ME), n=3. **B.** Non-reducing SDS-PAGE/Western blot analysis of failed FKBP2 KO (Ctrl 1 and 2) and FKBP2 KO cells cultured for 3 h in 20 and 2 mM glucose conditions. The presence of high molecular weight proinsulin complexes was evaluated with proinsulin specific antibody as in Arunagiri et al. [1], n=10. **C.** Void fractions of size exclusion chromatography of failed FKBP2 KO (Ctrl) and FKBP2 KO cells (cultured for 3h in 20 mM glucose containing media) were analysed under non-reducing and reducing conditions with SDS-PAGE/Western blot to detect proinsulin, n=3. **D.** Insoluble fractions of failed FKBP2 KO (ctrl 1 and 2) and FKBP2 KO cells cultured for 3 h in 20 and 2 mM glucose conditions were treated with 2% SDS containing loading buffer, boiled for 20 minutes and analysed by SDS-PAGE/Western blot to detect proinsulin, n=10. Proinsulin detection was done with CCI-17 monoclonal antibody. All Western blot quantifications were carried out with ImageJ. The data was analysed by the unpaired student t-test and is presented as means SD ±. **E.** Schematic representation of the proinsulin isomers folding steps proposed in this work.

We reasoned that some PI multimers in FKBP2 KO cells reach a state of lower solubility and consequently are not detectable in soluble cell fractions, resembling insulin-like growth factor 2 oligomerization that can transition into irreversible aggregates [37]. We therefore separated non-reduced control and FKBP2 KO cell lysates via SEC and analyzed its void fraction of protein aggregates. Upon SDS-PAGE and WB analysis, FKBP2 KO cells demonstrated a 3-fold increase in PI complexes as compared to controls (Fig. 6C). Finally, SDS-PAGE and WB analysis of cell pellets containing non-soluble proteins after cell lysis, demonstrated an over 40 % increase in the PI content in FKBP2 KO compared to control cells (Fig. 6D). These results point to a progressive accumulation of PI multimers with a subsequent loss of solubility due to impaired PI folding when cells lack functional FKBP2.

## Discussion

The β-cell intrinsic mechanisms underlying deficient insulin secretion in T2D are still unclear and much needs to be uncovered regarding PI biosynthesis, folding and the role of ER protein chaperones in this process. PI folding i.e. locking its 3D structure, is orchestrated in the ER, but apart from chaperoning by GRP94 and disulphide bridge formation, our insights into this process is limited [5, 6, 43]. This is in stark contrast to the recent identification of 38 PI-interacting proteins suggesting that multiple folding steps are required even before PI exits the ER and undergoes proteolytic cleavage to produce mature insulin [44]. Here, we discovered a novel role for FKBP2, a proline isomerase, in the folding of wild-type PI (Fig. 6E). We found that FKBP2 physically interacts with PI and the PI chaperone, GRP94 and that PI proline at position 28 is a likely target for FKBP2-dependent isomerization. Further we demonstrate that β-cells with FKBP2 KO exhibit loss of intracellular PI associated with increased misfolding, loss of solubility and increased PI secretion. Furthermore, FKBP2 KO cells showed increased apoptosis levels with no detectable changes in expression of ER stress markers. Finally, we observed FKBP2 overexpression in β-cells from human islets obtained from T2D patients, likely as a compensatory response to the increased insulin biosynthetic demand.

*In silico* prediction models of the PI-FKBP2 interaction suggested that proinsulin interacts with FKBP2 through a set of conserved residues [45, 46] involved in FKBP2’s binding of the inhibitor FK506, potentially necessary to facilitate proline isomerization (Fig. 2D). This suggested to us that FKBP2 might be a *bona fide* PI isomerase. Human PI contains three proline residues, one in the B-chain and two in the C-peptide; with only P28 in the B-chain, being conserved among species (Fig. 2E-F). Previous studies have shown that modifications of P28 have little to no effect on PI bioactivity, although they diminish PI dimer-formation [47, 48]. Somewhat contradictory to that, recently a P28L mutation was identified in a maturity-onset diabetes of the young (MODY) patient, and molecular analysis identified partial defects in PI (P28L) oxidative folding and ER export [20]. The discrepancy may be explained by the fact that initial P28 modifications or substitutions were introduced outside of the physiological folding environment that is found in β-cells whereas the P28L mutant undergoes folding attempts in vivo. In other words, determinants of foldability i.e. P28, may not be apparent once the native protein state is reached by other means i.e. *in vitro* oxidative PI folding [49, 50]. If correct, it may suggest that P28 is indispensable for proper PI folding during the transition through ER. The ability of proline to adopt a *cis* or *trans* configuration accounts for proline’s tendency to bend the regional amino acid arrangement, disturb protein secondary structure by inhibiting an α-helix or β-sheet conformation, therefore allowing for subsequent protein folding [51, 52]. In fact, FKBP2 binds more efficiently to unfolded PI than it’s folded state (Fig. 6A), indicating that the potential proline isomerization takes place before the PI structure is locked by the formation of disulphide bonds. Finally, FKBP2 in an *in vitro* isomerization assay was efficient in the *cis-*to-*trans* isomerization of proline residue in a nanopeptide derived from the human PI sequence directly preceding and containing P28 residue (Fig. 2G).

If FKBP2 works *in vivo* as a PI isomerase i.e. converts *cis* isomers to *trans*, and if *cis* isomers disturb or even prevent PI folding, we would expect PI to misfold in FKBP2 deficient cells. We thus knocked out FKBP2 (Suppl. Fig. 3) and demonstrated a consistent phenotype of lower cellular contents of PI (at 2 and 20 mM glucose concentrations) and insulin (at 20 mM; Fig. 3A). The PI content was partially restored after FKBP2 reconstitution in FKBP2 KO cells (Fig. 3B).

The β-cell has at least two ways of dealing with misfolded PI, by an increase in PI secretion and via intracellular PI degradation. In fact, if PI is not processed within 1 h to mature insulin, it is directed to and secreted via the constitutive pathway [53], and an elevated PI secretion is indeed a hallmark of T2D and T1D [54–56]. PI secretion and intracellular degradation result in a shorter PI half-life and in fact, this was our observation upon FKBP2 KO. As shown in Fig. 5B, shorter PI half-life in FKBP2 KO cells is partially a result of increased PI secretion, as prolongation of PI half-life was achieved via the treatment of cells with the ER-Golgi anterograde transport inhibitor, BFA. Importantly, FKBP2 KO impacts mostly PI folding status, as FKBP2 deficient cells maintain normal insulin secretion in response to glucose (Fig. 5D).

The only partial restoration of PI content in secretion-inhibited FKBP2 KO cells may suggest that misfolded PI is actively degraded as well. As mentioned, FKBP2 has a preference for binding unfolded, fully reduced PI. This would place the P28 isomerization before the formation of PI disulfide bonds. As a consequence, increased formation of aberrant PI disulphide bonds would be anticipated. Indeed, FKBP2 KO cells contain relatively more PI multimers, including high molecular weight complexes (HMWC) that have been reported to represent PI disulphide-linked multimers (Fig. 6B and Suppl. Fig. 4; [1]) where disulfide bonds are formed between different PI molecules. We supplemented this observation with the analysis of void SEC fractions, as large molecules (or their covalently linked complexes) have no or limited affinity to the SEC column resin pores and are thus eluted in the void fraction [57]. We found similar enrichment of PI disulfide-linked multimers in FKBP2 KO cells (Fig. 6C upper blot) that could be reduced (Fig. 6C lower blot) indicating that indeed these are multiple PI molecules linked via abnormal disulfide bonds. Furthermore, FKBP2 KO results in the accumulation of PI molecules in non-soluble cellular fractions (pellets, Fig. 6D). This may be a result of two factors: 1. Misfolded proteins, if not degraded promptly, tend to form aggregates i.e. insoluble high molecular weight forms [58]; 2. it has been suggested that the flexibility of the PI B-chain C-terminus (residues B27-B30) contributes to aggregation of insulin and formation of fibrils [59–61]. Consequently, modification to residues B27-B30 could lower PI solubility. Through the KO of FKBP2, we could potentially increase the presence of differentially spatially oriented and less soluble cis isomers. Taken together with the increase in HMWC, this may explain the accumulation of PI in the insoluble cellular fraction.

The above FKBP2 KO phenotype may explain why we did not observe changes to PI dimer formation (Fig. 3A). It is plausible that only PI molecules containing *trans* P28 are competent to form dimers and they represent the majority of detectable PI, while *cis* P28 isomers are secreted and/or misfolded and form non-soluble aggregates (Fig. 3C and 6D).

Models with PI misfolding in the ER display β-cell stress and apoptosis, insulin deficiency and the onset of diabetes [62]. In contrast, FKBP2 KO in β-cells led to the specific loss of PI and low-grade apoptosis but no detectable ER stress induction or other detrimental effects were observed (Fig. 4). How do we consolidate this induction of apoptosis without the activation of the ER stress pathways?

First, FKBP2 deficiency itself or through PI misfolding could potentially increases ER-to-mitochondria Ca^2+^ transport via the ER-mitochondria contact sites [63] and/or sustains an increase of cytosolic Ca^2+^, both inducing the intrinsic apoptotic pathway (reviewed in [64]), as is seen in prion-related disorders [65]. Second, the observed apoptosis may not be related to PI misfolding but rather represent a dysfunction of another FKBP2 target. In a recent survey of FKBP2 interacting proteins (data not shown), we identified subunits of mitochondrial ATP synthase that catalyzes ATP synthesis [66], or 60 kDa heat shock protein, HSPD1, which participates in the correct folding of proteins imported to mitochondria [67]. Interfering with the function of these proteins, could induce the intrinsic mitochondrial apoptotic pathway without inducing ER stress [68]. However, it remains to be investigated, if those proteins are functional FKBP2 targets. Third, during the generation of FKBP2 KO, cells that responded to KO and subsequent PI misfolding with ER stress followed by unresolved unfolded protein response, might not have survived the clonal selection. Hence, the FKBP2 KO clones that survived and were used during this work might have undergone adaptation, the result of which is tuning down ER stress and low levels of apoptosis.

Taken together, these results indicate a functional link between FKBP2 and PI folding. Under condition where an increased production of insulin is required (e.g. T2D), induction of FKBP2 expression may function as a compensatory response. Indeed, single-cell sequencing of pancreatic islets from human T2D donors demonstrated an up-regulation of FKBP2 mRNA in β-cells (Fig. 1E), however these data have not been independently validated (e.g. by qPCR).

In summary, our study has for the first time established the importance of FKBP2 as an essential PI isomerase in pancreatic β-cells and highlights a novel key function for PI P28 isomerization during PI folding. As such, our findings create an opportunity to investigate the potential of PI P28 isomerization status as a biomarker of early diabetes as well as a potential novel therapeutic target capable of improving PI folding.

## Abbreviations

ATF3/6: Activating Transcription Factor 3 and 6
BFA: Brefeldin A
BiP: Binding immunoglobulin Protein
CHX: Cycloheximide
CHOP: C/EBP homologous Protein
CRISPR/Cas9: Clustered Regularly Interspaced Short Palindromic Repeats/CRISPR-associated protein 9
Ctrl: Control
ER: Endoplasmic Reticulum
ERSE: ER Stress Response Element Site
FKBP2: FK506-Binding Protein 2
GFP: Green Fluorescent Protein
GRP94: Glucose Regulated Protein 94
GSIS: Glucose-Stimulated Insulin Secretion
HMWC: High Molecular Weight Complexes
HSPD1: Heat Shock Protein Family D Member 1
IRE1: Serine/threonine-protein inase/endoribonuclease Inositol-Requiring Enzyme 1
KO: Knock Out
KRBH: Krebs-Ringer’s-Bicarbonate-Hepes
MODY: Maturity-Onset Diabetes of the Young
PERK: Protein Kinase R-like Endoplasmic Reticulum Kinase
PI: Proinsulin
Pro: Proline
P28: Position 28 proline at of the proinsulin β-chain
SEC: Size Exclusion Chromatography
SP1: Specificity Protein 1
T2D: Type 2 Diabetes
XBP1: X-box Binding Protein 1

## Acknowledgements

We want to thank C. Wollheim and P. Maechler, University Medical Centre, Geneva, Switzerland for sharing with us the rat insulinoma INS-1E cell line.

## Funding

This project was funded by the EFSD/Lilly Programme, Dagmar Marshall Fond, Else og Mogens Wedell-Wedellsborgs Fond, Lægeforeningens Forsk, Magda Sofie og Aase Lütz Mindelegat, Augustinus Foundation (MTM) and National Science Center, Poland (MB), The Punjab Educational Endowment Fund (MSK). Collaboration with PEM was supported by a Visiting Professorship from the Danish Diabetes Academy (NNF17SA0031406).

## Conflict of interest

No conflict of interest is declared by any authors.

## Author contributions

CH, THB and CP developed the protocols for the experiments, conducted experiments, performed the statistical analysis, constructed figures and tables and wrote the manuscript. SNT (Fig. 3, 5, 6), SEB (Fig. 1, 2), HAH (Fig. 3, 6), MSK (Fig. 3, 6), KK (Fig. 1, 2, 5), MM and JD (Fig. 3, 5), EU and MB (Fig. 1) developed the protocols and conducted experiments, participated in critical data analysis and writing manuscript. TDS and PEM performed data analysis (Fig. 1). PMH planned, performed MS experiment and analyzed the data (Fig. 1). MP, PP and MW participated in the planning and evaluation of experiments related to the FKBP2-dependent PI isomerization (Fig. 2). PG and KG planned and supervised and analyzed experiments (Fig. 3, 6). TMPO, MJP, JK and RB participated in critical data analysis and writing of the manuscript. MTM is the study guarantor and was its initiator and leading supervisor; developed the protocols for the experiments, conducted experiments, performed the statistical analysis and constructed figures and tables as well as wrote the manuscript.

---

**S1.**
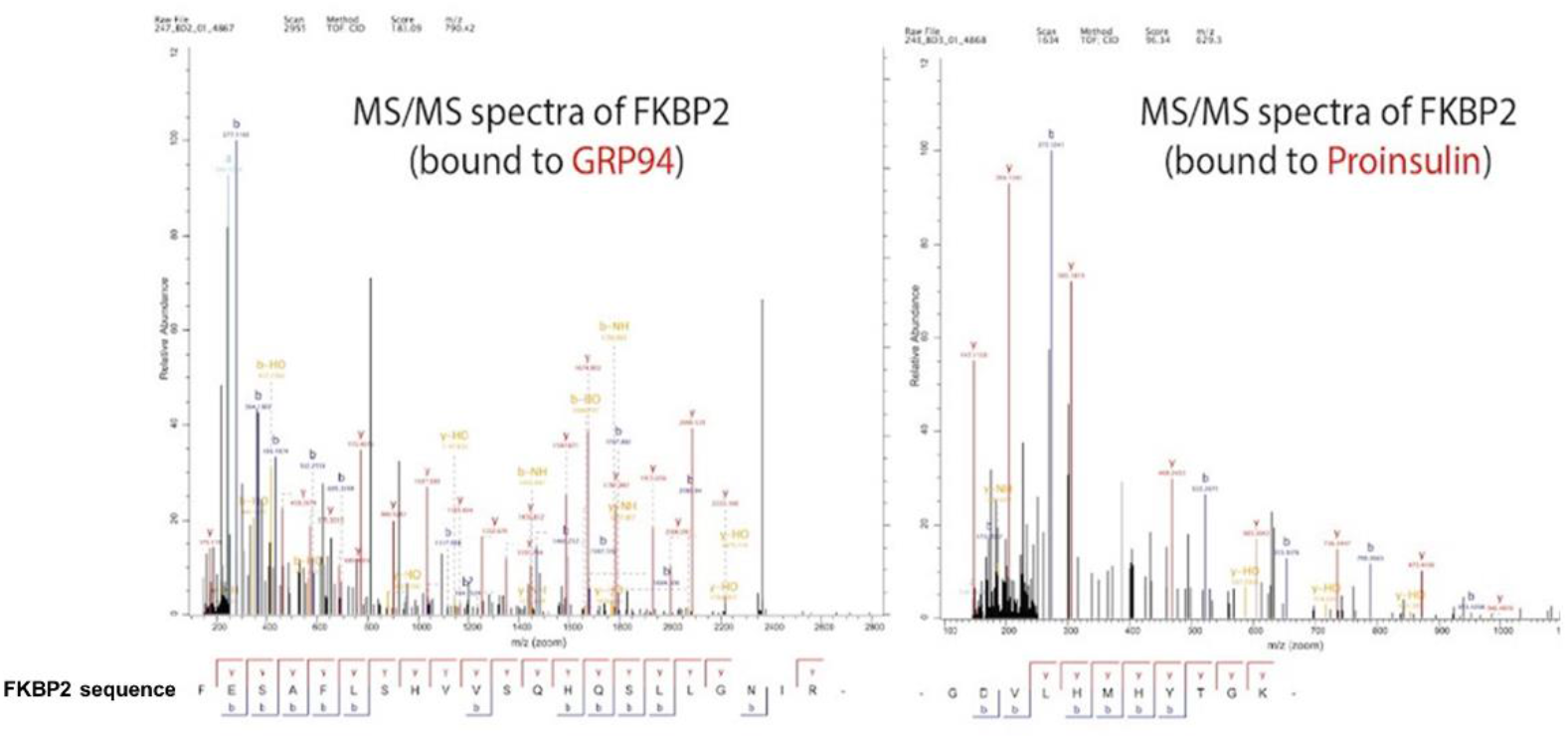
INS-1E cells were transiently transfected with GFP-taggcd GRP94 or GFP-tagged proinsulin, followed by GFP-dependent immunoprecipitation by GFP-tag and analysis via mass spectrometry (MS). MS/MS spectra of FKBP2 detected in GRP94 and proinsulin precipitates are presented.

**S2.**
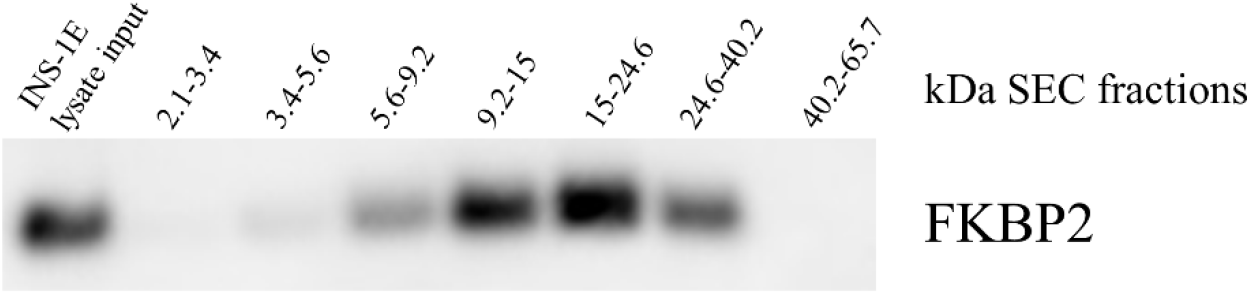
Size exclusion chromatography fractions with FKBP2 elution al molecular weights demonstrating that FKBP2 is expressed as a monomer in INS-1E cells.

**S3.**
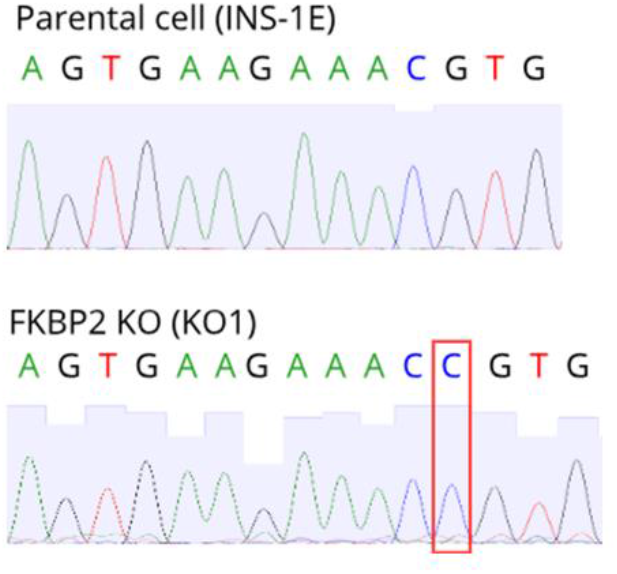
Sequencing results of exon 3 of *fkbp2* after CRISPR/Cas9 induced insertion of 1 bp C (indicated by red box) in FKBP2 KO clone 1 of INS-1E cells and its wild type counterpart in control clone.

**S4.**
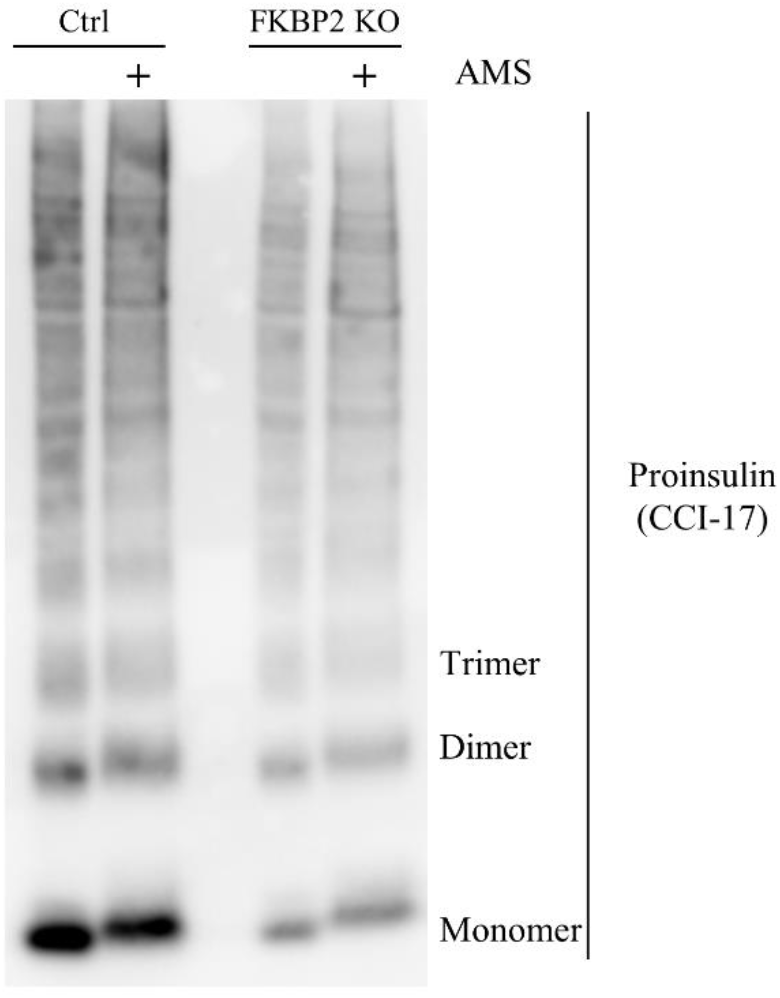
Detection of improperly folded proinsulin. INS1-E (Ctrl) and FKBP2 KO cells were lysed, divided into two portions with one alkylated in the presence of 6 mM 4-acetamido-4’-maleimidyl-stilbene2,2’-disulfonate (AMS) for 1 h, and then both resolved by non-reducing SDS-PAGE and analyzed by Western blotting with anti-proinsulin CCI-17 antibody from Novus Biologicals.

## Notes

### Competing Interest Statement

The authors have declared no competing interest.

