## Supplementary materials for "FK506-binding protein 2 participates in proinsulin folding"

**The Supplement contains:**

Supplementary materials and procedures

Supplementary Table 1- 4

Supplementary Figure 1 – 4

References

Original Western blots

**Cell culture**

The rat insulinoma INS-1E cell line was gifted by C. Wollheim and P. Maechler, University Medical Centre, Geneva, Switzerland. The wild-type and FKBP2 KO INS-1E cell lines were grown in RPMI-1640 GlutaMAX medium (11 mM glucose) supplemented with 10 % fetal bovine serum, 100 U/mL penicillin, 100 µg/mL streptomycin, 10 mM HEPES, 1 mM sodium pyruvate and 50 µmol 2-mercaptoethanol. Cells were incubated at 37˚C in a humidified atmosphere with 5 % $CO_2$. The medium was changed twice a week, and when the cells obtained a confluence of 80-90 %, they were split.

**Cell transfections**

Cells were transfected at 60-80 % confluence using Lipofectamine™ 3000 Transfection Reagent (ThermoFisher Scientific, Denmark) according to the manufacturer's protocol. Briefly, Lipofectamine™ 3000 reagent and Opti-MEM medium were mixed. In a separate tube, DNA (2.5 µg per $1x10^6$ cells) was diluted in Opti-MEM Medium to which P3000 reagent was added. The DNA-P3000 mix was mixed with the diluted Lipofectamine™ 3000 reagent. After 15 min incubation, the DNA-lipid complex was added to the cells dropwise and incubated for 16 hours. The next day transfection medium was replaced with regular growth media.

**Supplementary Table 1 Plasmids employed for exogenous protein expression**

| | | |
|---|---|---|
| Myc-tagged FKBP2 pcDNA3.1(+) | U0225EE200-2/M71546 | GeneScript, Netherlands |
| hProinsulinFLAGCpeptide pcDNA3.1(+) | U5277FA070-2/F11600 | GeneScript, Netherlands |
| cGRP94-GFP-KDEL | | [1, 2] |

| | | |
|---|---|---|
| hPro-CpepSfGFP | | [3] |
| GFP-KDEL (ER-localized GFP) | | [1, 2] |

**Immunoprecipitation and mass spectrometry analysis**

When $5x10^6$ INS-1E cells plated in a T75 flask reached ~70 % confluence, they were transfected with Myc-tagged FKBP2, cGRP94-GFP-KDEL, hPro-CpepSfGFP or GFP-KDEL using Lipofectamine™ 3000 (as described above). 48 hours post transfection, transfected cells were lysed in lysis buffer containing: 10 mM Tris pH 8, 150 mM Sodium Chloride, 0.5 mM EDTA, 2 % IGEPAL and protease inhibitor (Thermo Fisher Scientific, Denmark). GRP94, proinsulin and ER-localized GFP were immunoprecipitated with the magnetic GFP-trap beads (Chromotek, Germany). Beads were incubated with 8000 µg of protein from lysates (measured by Bradford assay, see later) for 1 hour, followed by six washes in dilution buffer (10 mM Tris pH 8, 150 mM sodium chloride and 0.5 mM EDTA). IP samples were reconstituted in 50 µL 8 M urea in 100 mM Tris/HCl pH 8.0, added 5 µL 100 mM DTT followed by incubation 30 min at room temperature, added 5 µL 200 mM iodoacetamide followed by incubation 30 min at room temperature in the dark. Subsequently 1 µL LysC (0.2 µg µL−1; Promega, Denmark) was added and samples were incubated for 3 h at room temperature, followed by addition of 250 µL 22 mM Tris/HCl pH 8.0 and 0.1 µL trypsin (1 µg µL−1; Promega, Denmark). Following incubation at room temperature overnight, samples were subjected to solid-phase extraction using StageTip C18 reverse-phase discs packed into pipette tips as described previously [4]. Samples were reconstituted with 20 µL 0.1 % formic acid and analyzed on a Bruker Impact II ESI-QTOF (Bruker Daltonics, Germany) mass spectrometer with an on-line Dionex Ultimate 3000 chromatography system (Thermo Fisher Scientific, Denmark) equipped with an Acclaim Pepmap C18 Column (15 cm, 300 µm ID). Peptides were eluted using a solvent gradient over 55 min, using acetonitrile with 0.1 % formic acid at a flow rate of 5 $\mu L\ min^{-1}$. Database searches were performed with MaxQuant v1.6.1.0 using the following parameters: enzyme: trypsin, with two missed cleavages; variable modifications: methionine oxidation; fixed modification: carbamidomethyl (cysteine); 1 % PSM and protein false discovery rate; mass tolerance: 0.07 and 0.005 Da (first and main searches, respectively); MS/MS mass tolerance: 40 ppm (first and main searches). Half of Myc-tagged FKBP2-containing cell lysates were pretreated for 10 minutes with 100 mM of reducing agent 2-mercaptoethanol prior to immunoprecipitation.

**FKBP2 expression in pancreatic endocrine cells**

Human pancreas sections were obtained from the Department of Clinical Pathology, Collegium Medicum in Bydgoszcz and Nicolaus Copernicus University in Torun (Poland) under protocols approved by the Bioethical Commission (KB381/2020). The donor identities were encrypted, and the data were analyzed anonymously. Only material with confirmed normal histology was included in the study. The pancreas was fixed in 10 % neutral buffered formalin for 24 hours (Alpinus Chemia, Poland), and further processed using the automated histo-processor (Excelsior AS; Thermo Fisher Scientific, Poland), according to routine protocol. The 4 μm sections of adhered glass slides (SuperFrost plus, Bionovo, Poland) were deparaffinized and subjected to a citrate buffer antigen retrieval protocol before proceeding to immunofluorescence staining (IF). The samples were then incubated with primary antibodies diluted in NDS overnight at 4° C. Primary antibodies used in the study are listed in Supp. Table 2. After two 10-min washes with 0.1 % Tween-20 in Phosphate-buffered saline (PBS), cells were incubated with secondary antibodies conjugated to AlexaFluor488, TRITC or AlexaFluor647 (all Jackson Immunoresearch, USA) diluted 1:800 in NDS for 2 hours at room temperature. Secondary antibodies were then washed out twice for 5 min with PBS-Tween-20 and incubated with DAPI (Sigma Aldrich, Poland) as a counterstain. Tissue samples were mounted with ProLong Diamond Antifade (Thermo Fisher Scientific, Poland) prior to imaging with Nikon Confocal and Leica inverted microscope DMIL LED.

**Supplementary Table 2 Antibodies for immunostaining**

| Antibody | Catalog number | Company |
|---|---|---|
| FKBP2 | sc-390753 | Santa Cruz |
| Insulin | 3014S | Cell Signaling |
| Somatostatin | SC-7819 | Cell Signaling |

#### Immunoblotting

**Cell lysates** The cells were washed in Hank's Balanced Saline Solution (HBSS, (without $Ca^{2+}$ and $MG^{2+}$) and lysed in lysis buffer (10 mM Tris pH 7.4, 150 mM NaCl, 0.1 % SDS, 1 % NP40, 2 mM EDTA, 1M $ZnCl_2$, supplemented with protease inhibitor (ThermoFisher Scientific, Denmark). After 30 minutes of incubation on ice, lysates were centrifuged at 10,000 x g for 25 minutes at 4˚C and protein concentration was determined using the Protein Assay Dye Reagent Concentrate (Bio-Rad, Denmark). After adjusting for protein concentration, samples were prepared in lysis buffer and NuPAGE LDS sample buffer (Bio-Rad, Denmark) with or without 10 % 2-mercaptoethanol (Sigma Aldrich, Denmark). Samples were heated to 95˚C for 5 min and briefly spun down. 5-30 µg of protein and protein marker (Bio-Rad, Denmark) were separated under reducing or non-reducing conditions on Nu-Page 4-12 % Bis-Tris Protein Gels (ThermoFisher Scientific, Denmark) at 150 V for 65 minutes and transferred to PVDF membranes by electro-transfer. Upon transfer, the membrane was rinsed in MilliQ water, blocked in 1 % non-fat milk in TBS (15 mM Tris, 150 mM NaCl) for 1 hour, and washed 3x10 min in TBST (15 mM Tris, 150 mM NaCl, 0.1 % Tween-20). Primary antibodies were diluted in 2 % BSA TBST and incubated with membranes overnight at 4˚C. The membranes were washed 3x10 min in TBST prior to 1 hour blotting with secondary antibody. Secondary HRP-conjugated antibodies were diluted 1:10.000 in 1 % non-fat milk in TBST and washed for another 3x10 minutes in TBST. Blots were developed using a chemiluminescence detection system (Radiance Q Chemiluminescent Substrate for quantitative Westerns, Azure Biosystems, USA; Clarity$^{TM}$ Western ECL, Bio-Rad, USA; SuperSignal West Femto Maximum Sensitivity Substrate, Thermo Scientific, Denmark), and light emission was captured using the Azure Sapphire Biomolecular Imager system (Azure Biosystems Inc, USA). For quantification of the protein bands, ImageJ 1.53e software was used.

**AMS-mediated alkylation of proinsulin** Alkylation of proinsulin was determined as recently described in [5]. Briefly, INS-1E wild-type and FKBP2 KO cells were prepared in reaction buffer (50mM Tris, pH 7.4) and NuPAGE LDS sample buffer (Bio-Rad, Denmark). Samples were heated to 95˚C for 10 min and cooled to 37˚C. Upon cool-down, 6 mM 4-Acetamido-4'-Maleimidylstilbene-2,2'-Disulfonic Acid, Disodium Salt (ThermoFisher Scientific, Denmark) was added to the samples, which were vortexed, spun down, and incubated further for 1 hour at 37˚C. 5-10 µg protein of the AMS-treated and untreated controls were loaded onto a non-reducing Nu-Page 4-12 % Bis-Tris Protein gel, which was incubated in 25 mM Dithiothreitol (DTT) for 10 minutes at room temperature prior to electro-transfer to a PVDF membrane.

**Proinsulin half-life** was evaluated by inhibition of protein synthesis with cycloheximide (CHX). CHX was purchased as a ready-to-use solution (cat. #C4859, Sigma Aldrich, Denmark) and used at 100 µM. Brefeldin A (cat. #B6542, Sigma Aldrich, Denmark), an inhibitor of ER-to-Golgi transport and consequently protein exocytosis, was reconstituted with DMSO and used at a concentration of 200 nM.

**Cellular pellets** Upon cell lysis, the supernatant (lysate) was removed and kept for further analysis. The remaining pellets (insoluble cellular remnants) were mixed with 17 µL of lysis buffer and 3 µL of NuPAGE sodium dodecyl sulphate (SDS) sample buffer (Bio-Rad, Denmark) containing 2-mercaptoethanol and then homogenized with a sharp needle attached to a syringe. The samples were heated at 95°C for 20 minutes and thereafter run on a Nu-Page 4-12 % Bis-Tris Protein Gel and subsequently transferred to PVDF membranes by electro-transfer and processed as mentioned above.

**Cell culture supernatants** One million cells were plated on a 6-well plate and incubated at 37°C, 5 % $CO_2$ and 95 % humidity. After 48 hours the old supernatant was discarded and exchanged for 1 mL of fresh medium. After 3 hours, the supernatant was collected and transferred to precooled Eppendorf tubes. The supernatants were then concentrated according to protein size, using the Vivacon® 500 centrifugal units (Sartorius-Fisher Scientific, Denmark). 500 µL of the supernatant was transferred three times to the hydrostat membrane and the assembled concentrator was centrifuged at 10.000 g for 3 x 30 minutes. 6.5 µL (or adjusted volume according to cell count) of the concentrated supernatant was suspended into an Eppendorf Tube together with 10 µL lysis buffer and 5.5 µL NuPAGE SDS sample buffer containing 2-mercaptoethanol. The solution was heated at 95°C for 5 minutes and run on a Nu-Page 4-12 % Bis-Tris Protein Gel and subsequently transferred to PVDF membranes by electro-transfer and processed as mentioned above.

**Supplementary Table 3 Primary and secondary antibodies used**

| Antibody | Catalog number | Company | Dilution |
|---|---|---|---|
| anti-insulin | 8138 | Cell Signaling, USA | 1:1000 |
| anti-proinsulin | NB100-73013 | Novus Biologicals, Denmark | 1:1000 |
| anti-FKBP2 | sc-390753 | Santa Cruz Biotechnology, USA | 1:1000 |
| anti-GRP94 | ADI-SPA-850 | Enzo Life Sciences, Denmark | 1:5000 |
| anti-tubulin | T6074 | Sigma Aldrich, Denmark | 1:2000 |
| anti-GAPDH | ab8245 | Abcam, Great Britain | 1:2000 |
| anti-rat | ab6734 | Abcam, Great Britain | 1:10000 |
| anti-rabbit | 7074S | Cell Signaling, USA | 1:10000 |
| anti-mouse | 7076S | Cell Signaling, USA | 1:10000 |

**FKBP2 protein activity assay**

n-Nitroanilide modified peptide (GERGFFYTPF-F-pNA, U1529Ge060-1, GenScript, USA) was mixed with assay buffer (PBS pH 7.4) to a final concentration of 5 mM, 170 µL of the solution was added to a flat bottom 96 well plate. 10 µL (1 µg) of recombinant FKBP2 (ab93681, Abcam, Great Britain) in assay buffer or assay buffer alone (negative control) and 5 µL (20 nM) of chymotrypsin (Roche, Switzerland) were added and mixed by shaking before the readout. A mixture of assay buffer, peptide, and FKBP2 was used as blank. The increase in absorbance (at 405 nm) was read at 1 min intervals for 12 hours at 25°C. Signals from all reactions were collected at the time point where a reaction containing FKBP2, n-Nitroanilide modified peptide and chymotrypsin was the highest.

**Generation of FKBP2 CRISPR/Cas-9 mediated knockout INS-1E cell lines**

$5x10^4$ INS1-E cells were plated on fibronectin-covered 48-well plates. For transduction of cells, All-in-one Lentiviral sgRNA particles containing a mCMV promotor and the Cas9 nuclease with the guide RNA sequence GATTGGAGTGAAGAAACGTG (target ID VSGRNOM_29216786) targeting exon 2 of the rat *fkbp2* gene was used (Dharmacon, Denmark). When cells reached near full confluence, Lentiviral particles at either 5 or 0.5 MOI and fresh medium were added to a total volume of 500 µL. Fresh medium was supplied after 18 hours of transduction, and after an additional 24 hours, puromycin (Sigma Aldrich, Denmark) selection at 2 µg/mL was initiated. Cells were cloned by array dilution in fibronectin-covered 96-well plates in 200 µL fresh medium and allowed to propagate until clonal islets could be moved to larger culture plates also covered in fibronectin. FKBP2 KO was confirmed in 4 clones by Sanger sequencing of the rat *fkbp2* gene (Genewiz Inc., UK) and FKBP2 protein expression levels were evaluated by Western blotting. An additional four clones were selected where *fkbp2* targeting was unsuccessful (*fkbp2* wild-type sequence and FKBP2 protein level expression similar to those of parental INS-1E cells).

**qRT-PCR**

INS-1E cells ($1\times10^6$ per condition) were seeded in 6-well plates 48 hours before harvest. Total RNA was harvested and extracted using the Nucleo-Spin kit (Macherey-Nagel, Switzerland) according to the manufacturer's instructions. The quality and quantity of the extracted RNA were assessed using a NanoDrop 1000 (Thermo Fisher Scientific, Denmark). 500 ng of total RNA was used for cDNA synthesis with the iScript™ cDNA Synthesis Kit (BioRad, Denmark). Real-time qPCR was performed with 15 ng of cDNA with SYBR Green PCR Master Mix (Applied Biosystems from Thermo Fisher Scientific, Denmark) and specific primers (Suppl. Table 4). A total volume of 10 µL was loaded on a 384-well plate and run in a Real-Time PCR machine (QuantStudio 5, Thermo Fischer Scientific, Denmark) with thermal cycles as follows: hot start 95˚C, 10 min; 45X amplification 95˚C, 15 sec, 60,8 ˚C, 20 sec, 60˚C, 40 sec; dissociation curve 95˚C, 15 sec, 60˚C, 15 sec, 95˚C, 15 sec. Gene expression levels were normalized to 5S rRNA, H2A.Z Variant Histone 1 (H2az1) and RNA polymerase subunit 2 (RPII).

**Supplementary Table 4 Primers used during qRT-PCR**

| Primers | Forward | Reverse |
|---|---|---|
| 5S rRNA* | 5’ TCTTTGGGAAATGGAGCACT 3’ | 5’ ATGAGCTTCTTGCCGTTGTT 3’ |
| ATF6 | 5’ CTCAAAGTCCCAAGTCCAAAG 3’ | 5’ CACTCTCCTGGAATCTTGCTC 3’ |
| BiP | 5’CTGGCACTATTGCTGGACTG 3’ | 5’CCACCACTTCAAAGACAGCA 3’ |
| CHOP | 5’ CAGCGACAGAGCCAAAATACC 3’ | 5’ TGTGGTGGTGTATGAAGATGC 3’ |
| H2az1* | 5’ CGGTAAGGCTGGAAAGGAC 3’ | 5’ CGTGGCTTGTTGTCCTAGATT 3’ |
| INS1 | 5’ GGGGAACGTGGTTTCTTCTAC 3’ | 5’CCAGTTGGTAGAGGGAGCAG 3’ |
| INS2 | 5’CAGCACCTTTGTGGTTCTCA 3’ | 5’CACCTCCAGTGCCAAGGT 3 |
| IRE1 | 5’ TGCTCAAGGACATGGCTACTA 3’ | 5’ GAAGGAGCTGAATTTTCTCCA 3’ |
| Perk | 5’ ACAGCCAATGAGGAAGTTTTG 3’ | 5’ GTAGGGAACTTTTCCGAGACC 3’ |
| RPII* | 5’ GGAGAGGAGATGGACAACAAG 3’ | 5’ TCCATTCAGCATACAACTCCA 3’ |
| sXBP-1 | 5’CTGAGTCCGAATCAGGTGCAG 3’ | 5’ATCCATGGGAAGATGTTCTGG 3’ |
| usXBP-1 | 5’CAGCACTCAGACTACGTGCG 3’ | 5’ATCCATGGGAAGATGTTCTGG 3’ |

* rRNA (reference)

**Apoptosis Assay**

Apoptotic cell death was determined by the detection of DNA–histone complexes present in the cytoplasmic fraction of cells using the Cell Death Detection ELISAPLUS kit (Roche, Switzerland) as described by the manufacturer. Briefly, 1x$10^4$ INS-1E and FKBP2 KO cells were plated in 96-well plates. 0,7 µM of thapsigargin, added 18 hours before cell harvest, was used to induce ER stress. Twenty hours later, cells were spun down for 10 min at 200×g and lysed in 200 µL lysis buffer for 30 min at room temperature. Lysates were centrifuged for 10 min at 200×g, and 20 µL supernatant and 80 µL immunoreagent (anti-DNA–POD antibody and anti-histone–biotin antibody) were added to streptavidin-coated microtiter plates and incubated for 2 hours under shaking conditions (300 rev/min) at room temperature. The solution was then removed, and each well was washed three times with 250 µL incubation buffer, afterwards 100 µL ABTS solution was added. Absorbance was measured at 405 nm and 492 nm as reference.

**Size exclusion chromatography**

Cells were grown to 90 % confluence in T75 flasks and were incubated for 3 hours with fresh media containing 20 mM glucose. The number of cells harvested by trypsinization was adjusted by cell count. The cell pellets were washed in HBSS and lysed in standard lysis buffer. The lysates were incubated on ice for 30 minutes prior to centrifugation at 10,000 x g for 30 minutes at 4˚C and the soluble fractions were collected for further analysis. Protein concentration was determined using the Protein Assay Dye Reagent Concentrate (Bio-Rad, Denmark). The lysates were fractionated in Tris buffer (50 mM Tris pH 8.0, 150 mM NaCl, 5 mM KCl) using a Superdex™ 75 10/300 GL column (Sigma Aldrich, Denmark). Fractions of interest were collected and proceeded for SDS-PAGE and analyzed by immunoblotting.

**Glucose-stimulated insulin secretion**

$3x10^5$ INS-1E cells (control and FKBP2 KO) were seeded in 24-well plates in 2 mL complete RPMI medium. 48 hours later at 80 % confluence medium was removed, and cells were incubated for 2 hours in Krebs-Ringer's-bicarbonate-Hepes (KRBH) buffer (149 mM NaCl, 4.4 mM KCl, 1.32 mM $NaH_2PO_4(H_2O)$, 1.32 mM $MgCl_2$, 2.75 mM $CaCl_2$, 25 mM $NaHCO_3$,

10 mM HEPES pH 7.4 and 0.1 % BSA) with 2 mM glucose to establish basal insulin secretion. Cells were subsequently incubated with KRBH buffer containing 2 mM glucose for 1 hour, followed by 20 mM glucose for additional hour. Supernatants were collected for measurements of proinsulin and insulin (Mercodia, Sweden) according to the manufacturer's protocols. In short, the samples were diluted with calibrator 0 to 1:100-1000. 50 µL of enzyme conjugate 1X solution was added to each well. 25 µL of samples calibrators and controls were pipetted to the appropriate wells. After two hours incubation on a shaker (700-900 rpm) at room temperature, the plate was washed 6 times with 700 µL wash buffer 1X solution per well using an automatic plate washer. Then 200 µL of substrate TMB was added to each well. The plate was incubated for another 30 minute at room temperature (18-25$^0$C) before 50 µL stop solution was added to each well and incubated on a shaker for 5 seconds. Absorbance was measured at 405 nm.

### Original Western blots for all figures

Fig. 1A coIP

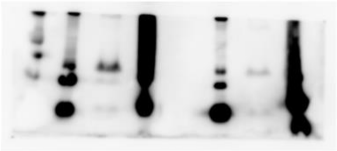

FKBP2

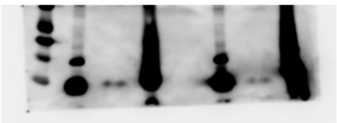

Proinsulin

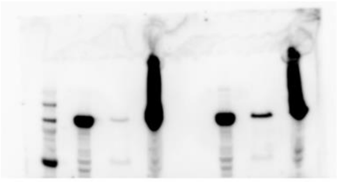

GRP94

Fig. 3A FKBP2 KO 2mM Glucose

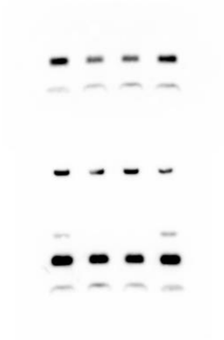

Proinsulin short exposure

Tubulin

Proinsulin long exposure

Fig. 3B FKBP2 reconstitution

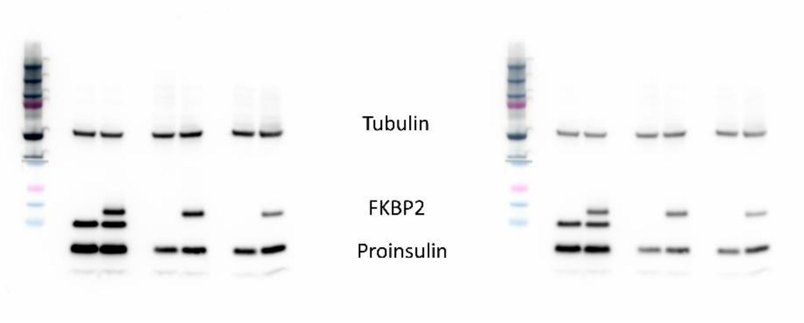


Fig. 3C SEC

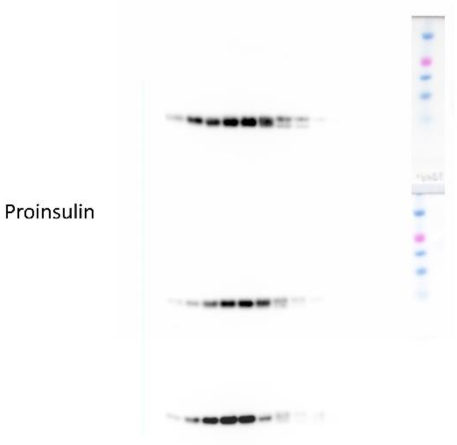


Fig. 5A Proinsulin half-life

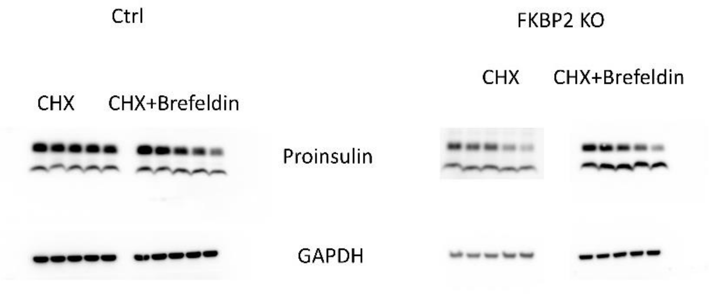

Fig. 5C Cell supernatants

Soups 20mM

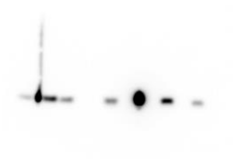

Soups 2mM

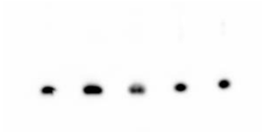

Fig. 6A coIP

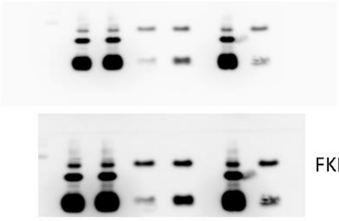


Fig. 6B

20mM CCI17

2mM CCI17

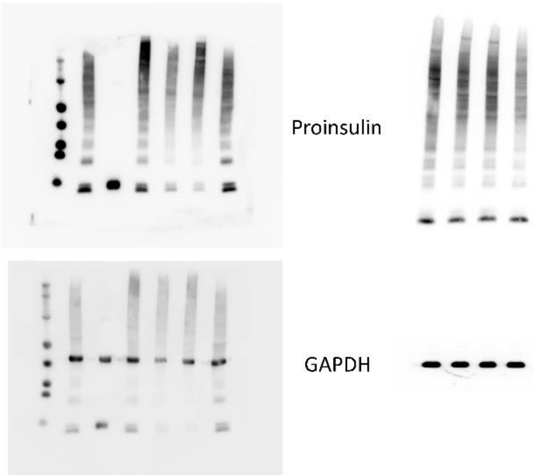

Fig. 6C void fractions

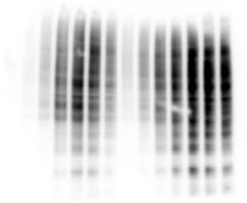

Proinsulin

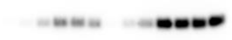

Proinsulin

Fig. 6D Cell pellets

Pellets 20mM

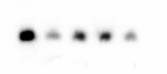

Pellets 2mM

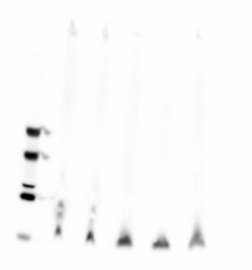

Suppl. Fig. 2 FKBP2 SEC distribution

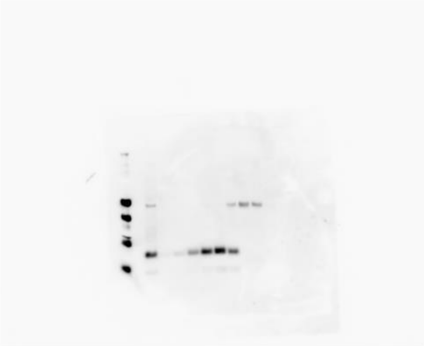

FKBP2

Suppl. Fig. 4 AMS treatment

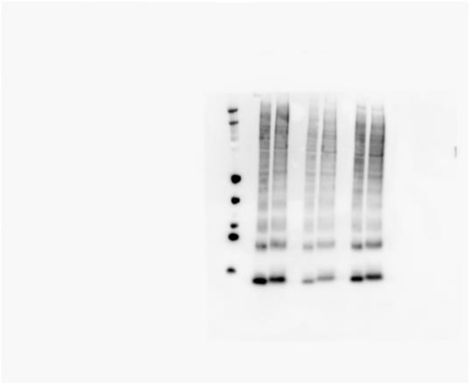
